# *Streptomyces* extracellular vesicles are a broad and permissive antimicrobial packaging and delivery system

**DOI:** 10.1101/2023.10.02.560488

**Authors:** Kirsten J. Meyer, Justin R. Nodwell

## Abstract

*Streptomyces* are the primary source of bioactive specialized metabolites used in research and medicine, including many antimicrobials. These are presumed to be secreted and function as freely soluble compounds. However, increasing evidence suggests extracellular vesicles are an alternative secretion system. We assessed environmental and lab-adapted *Streptomyces* (sporulating filamentous actinomycetes) and found a majority of strains produce vesicles packaging antimicrobial compounds. The cargo included actinomycins, anthracyclines, candicidin, and actinorhodin, reflecting both diverse chemical properties and diverse antibacterial and antifungal activity. The production of the antimicrobials and vesicles was linked, although vesicle production continued in the absence of antimicrobials. We demonstrated that antimicrobial-containing vesicles achieve direct delivery of the cargo to other microbes. Notably, this delivery via membrane fusion occurred to a broad range of microbes including pathogenic bacteria and yeast. Vesicle encapsulation offers a broad and permissive packaging and delivery system for antimicrobial specialized metabolites, with important implications for ecology and translation.

**GRAPHICAL ABSTRACT:** 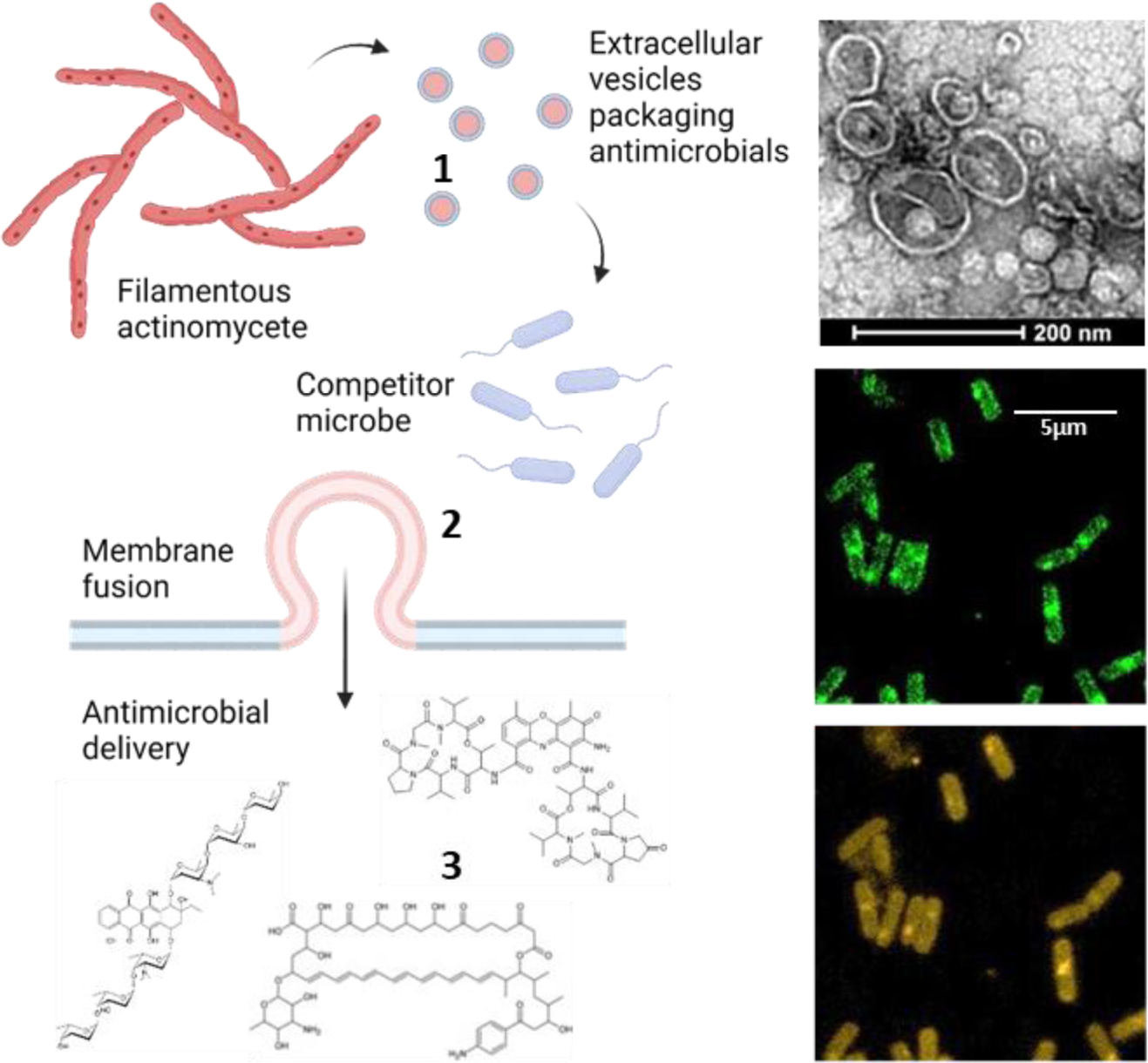

**IMPORTANCE:** Extracellular vesicle encapsulation changes our picture of how antimicrobial metabolites function in the environment, and provides an alternative translational approach for delivery of antimicrobials. We find the majority of *Streptomyces* strains are capable of releasing antimicrobial vesicles, and at least four distinct classes of compounds can be packaged, suggesting this is widespread in nature. This is a striking departure from the primary paradigm of secretion and action of specialized metabolites as soluble compounds. Importantly the vesicles deliver antimicrobial metabolites directly to other microbes via membrane fusion, including to pathogenic bacteria and yeast. This suggests future applications in which lipid-encapsulated natural product antibiotics and antifungals could be used to solve some of the most pressing problems in drug resistance.

## INTRODUCTION

Most of modern medicine would be unimaginable without microbial specialized metabolites. These molecules have potent biological activities and, as a result, have been developed as antibiotics, chemotherapeutics and other medicines^1^. The crisis of antimicrobial resistance and the relatively poor performance of synthetic chemical libraries in antimicrobial discovery has spurred a renaissance in specialized metabolite research^2^. Moreover, advances in genetic and metabolomic tools have demonstrated an abundance of novel chemistry yet to be explored in environmental bacteria^2–6^.

Early reports of clinically valuable microbial specialized metabolites came with the discovery of penicillin from the *Penicillium* mould in 1928 and streptomycin from *Streptomyces griseus* in 1943^7–9^. Since then, the majority of specialized metabolites have been mined from the enormous actinomycete bacterial genus *Streptomyces*^6^. Each strain of *Streptomyces* carries gene clusters for dozens of different specialized metabolites. Some of these compounds serve their producing organisms as siderophores and spore pigments^10^, and many act as antimicrobials^11^. Despite decades of research, mechanistic knowledge is limited to a relatively small fraction of these compounds, and is even more scarce at the level of endogenous production and function^12^.

Since their discovery nearly a century ago, it has been assumed that actinobacterial specialized metabolites generally act as free, soluble chemical agents and indeed, this is how they are employed clinically. The fact that many specialized metabolic gene clusters encode dedicated ABC transporters and efflux pumps has tacitly supported that view^13–15^. However, this view was challenged in 2011 when Schrempf and colleagues reported the ground-breaking discovery that actinorhodin, which had been studied for more than 60 years as a soluble compound^16,17^, was associated with lipid-encapsulated vesicles in secreted exudates from *Streptomyces coelicolor*^18^. Since then, antibiotic containing extracellular vesicles have been detected from *Streptomyces lividan*s, *Streptomyces* sp. Mg1 and *Streptomyces albus*^19–22^. These intriguing reports beg the question of how widespread this phenomenon might be amongst actinomyces.

It is increasingly apparent that extracellular vesicles are an integral part of the microbial world^23,24^. Studies have focused primarily on Gram-negative genera, where they are predominantly derived from the outer membrane and referred to as outer membrane vesicles (OMVs)^24–27^. Extracellular vesicles have now also been recognized in Gram-positive bacteria, forming from the cytoplasmic membrane by a poorly understood mechanism that is still under active investigation^28–30^. Vesicles are known to carry a variety of molecular cargo, with most reports focused on their protein content. Importantly, the specific set of proteins contained in these vesicles differs from that of the periplasm or cytoplasm of the bacteria, suggesting they might be loaded according to specific targeting or enrichment mechanisms^31–34^.

Many roles have been assigned to extracellular vesicles, ranging from the removal of macromolecular waste to the delivery of specific cargo^29^. Several pathogens deliver virulence factors such as α-hemolysin (*Staphylococcus aureus*)^35^ and cholera toxin (*Vibrio cholerae*)^36^ to host cells via extracellular vesicles. *Pseudomonas aeruginosa* releases vesicles that carry the quorum sensing molecule PQS^37^. Vesicles serve to facilitate horizontal gene transfer in *Acinetobacter baumannii*^38^. They can also serve as antagonistic agents^39^, carrying bacteriocins from *Lactobacilli*^40^, or lytic enzymes and toxic metabolites from *Myxococci*^41^, to kill other bacterial strains.

We asked what implications vesicle packaging might have for the transport and delivery of specialized metabolites, in particular antimicrobial metabolites. First, how widespread is this phenomenon in the most prolific specialized metabolite producers, the filamentous actinomycetes? Second, is vesicle production coincident with or catalyzed by the metabolites themselves? Third, and most importantly, how does vesicle encapsulation affect the delivery of specialized metabolites?

To address these questions, we examine eleven soil actinomycete isolates and six lab adapted strains for extracellular vesicle production and association with antimicrobial specialized metabolites. We find that under laboratory conditions this is indeed common, nine out of seventeen strains, for actinomycetes, including eight of eleven environmental isolates. We identify the antimicrobial specialized metabolites associated with extracellular vesicles from four strains, and find they have diverse chemical structures and biological activities. We demonstrate that vesicle production and antimicrobial production are temporally linked, and that the antimicrobial specialized metabolites are responsible for vesicle efficacy. Importantly, we find that these extracellular vesicles fuse with the membranes of diverse bacteria and fungi, delivering antimicrobials directly to the cells. Vesicle-mediated delivery as a broad, permissive, and common mechanism represents a new insight into the enormous family of antimicrobial specialized metabolites with far-reaching implications for their use in medicine and research.

## RESULTS AND DISCUSSION

### Filamentous actinomycetes frequently produce antimicrobial extracellular vesicles

If the packaging of specialized metabolites into extracellular vesicles is widespread, we reasoned it should be possible to detect metabolite-associated bioactivity in high molecular weight (MW) fractions of culture supernatants. Moreover, because vesicles are membrane-bounded, these high molecular weight fractions should also contain lipids. Using centrifugal size filtration, we created a rigorously washed high MW fraction >100 kDa and a matched fraction of low MW supernatant from one-week liquid cultures of seventeen antimicrobial producing actinomycetes. This included common lab-adapted strains and environmental soil isolates^42^. To test for lipids we added the lipid-reactive dye FM 4-64 to each fraction. Fourteen of seventeen high MW fractions had increased fluorescence compared to the low MW fractions, consistent with the presence of extracellular vesicles (Fig 1A, Supplementary Table 1).

**Figure 1.**
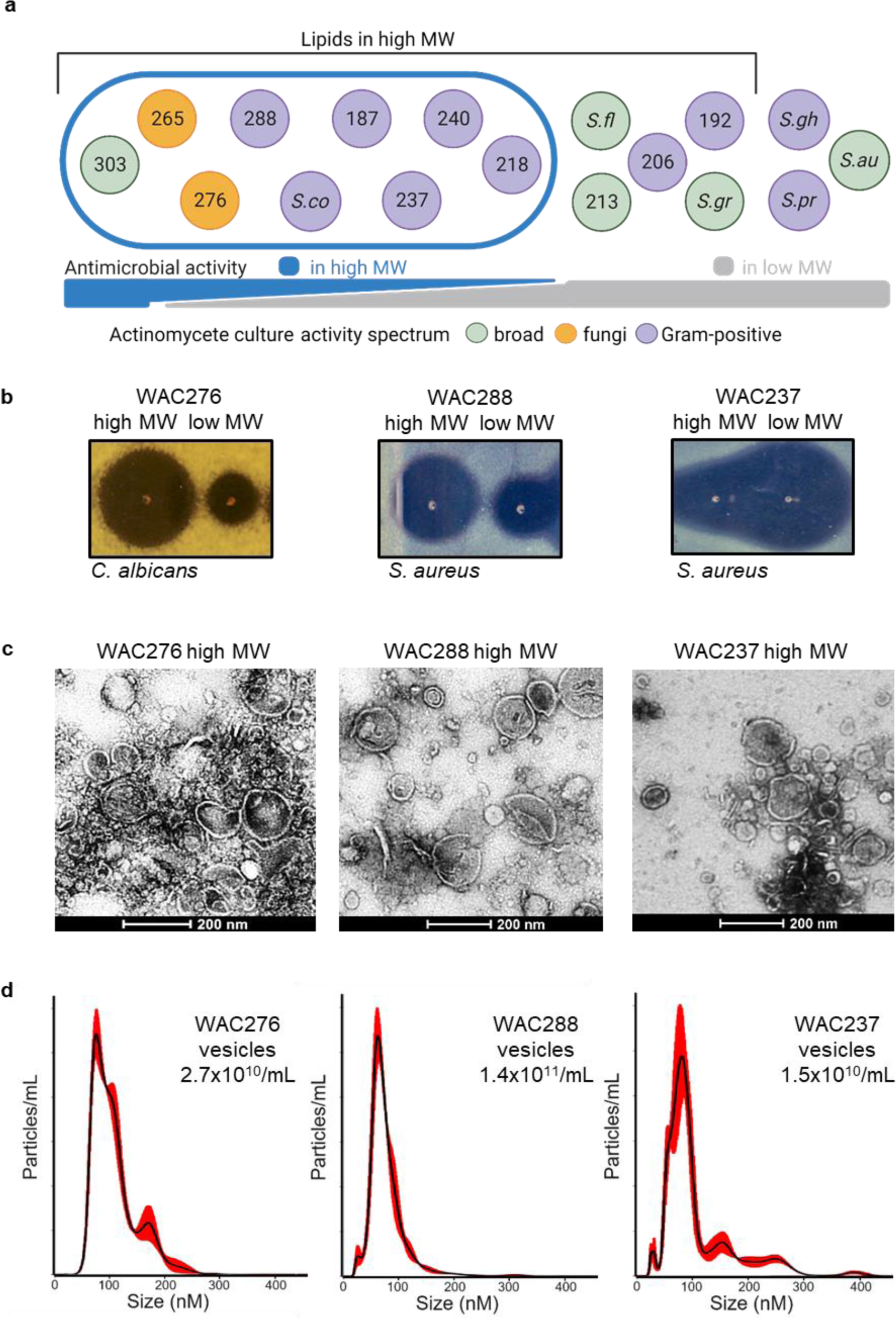
Filamentous actinomycetes commonly produce antimicrobial extracellular vesicles. (A) Seventeen actinomycete liquid culture supernatants were separated into a high molecular weight (MW) fraction >100 kDa and a low molecular weight fraction <100 kDa and tested for presence of lipids using lipid reactive dye FM 4-64 (fourteen out of seventeen high MW fractions positive, black bracket), and for antimicrobial activity. Activity was assessed against fungi *C. albicans* (yellow), Gram-positive bacteria *S. aureus* and *B. subtilis* (purple), or Gram-negative bacteria *K. pneumoniae*. Broad spectrum inhibition (more than one category) represented as green. Nine high MW fractions exhibited antimicrobial activity (encircled with blue), with the ratio of antimicrobial activity varying between the high MW fraction (blue) and the low MW fraction (grey), bottom bar. For example, (B), WAC276 high and low MW fractions created zones of inhibition against C. albicans, and WAC288 and WAC237 high and low MW fractions inhibited *S. aureus*. These high MW fractions were imaged by negative stain transmission electron microscopy revealing abundant extracellular vesicle structures (C). Size exclusion chromatography on culture supernatant followed by nanoparticle tracking analysis was used to measure the concentration per mL (total vesicle concentration top right inset) and the size distribution of these vesicles (D). Results are representative of at minimum biological duplicate.

Strikingly nine of these high MW fractions also had antimicrobial activity (Fig 1A, Supplementary Table 1). As expected for activity arising from specialized metabolites, the spectrum of microbial inhibition of the high MW fractions differed across isolates. Moreover, matching antimicrobial activity was present in the low MW fractions presumably due to a portion of metabolites free in solution (Fig 1B). Six high MW fractions, from the lab-adapted strain *S. coelicolor* M145, and the soil isolates WAC00187, 00218, 00237, 00240, and 00288, inhibited Gram-positive bacteria *Staphylococcus aureus* and *Bacillus subtilis.* Two high MW fractions, from WAC00265 and 00276, inhibited the fungi *Candida albicans*, and there was one, from WAC00303, with broad spectrum activity against both the bacteria and *C. albicans*. Examining the partitioning of antimicrobial activity between the high and low MW fractions, five of the isolates had equal or weaker activity in the high MW fraction (*S. coelicolor*, WAC00187, 00218, 00237, and 00240), whereas in four isolates (WAC00265, 00276, 00288, and 00303) the high MW fraction had the greatest antimicrobial activity (Fig 1B). Finally, to confirm extracellular vesicle production, we examined the high MW fractions from isolates across the range of antimicrobial profiles, *S. coelicolor* (Supplementary Fig 1A) and three of the environmental isolates, WAC00276, 00288, and 00237 (Fig 1C), using transmission electron microscopy. We observed abundant extracellular vesicles that ranged in size from tens to hundreds of nanometers in diameter (Fig 1C).

### Extracellular vesicles package diverse antimicrobial specialized metabolites

Taking four isolates with a range of bioactive profiles, we used LC-MS/MS to determine if the antimicrobial extracellular vesicles were associated with known specialized metabolites. The vesicle fraction from *Streptomyces coelicolor* contained a clear peak of the molecular ion expected for the specialized metabolite actinorhodin (Supplementary Fig 1A), corroborating previous work^18,43^. For the environmental isolates WAC00276, 00288, and 00237, we further purified the antimicrobial extracellular vesicles using size exclusion chromatography^44^. The concentration and size of vesicles was quantified using nanoparticle tracking analysis (Fig 1D). Vesicle concentrations were 10^10^ to 10^11^/mL of culture supernatant. Most vesicles were between 50 to 150 nm, with the median size falling between 60 to 100 nm. Extracts of the vesicles were run on LC-MS/MS, and comparison to blank controls and between the vesicles identified unique vesicle metabolite cargo. LC-MS/MS profiles of the bioactive HPLC fractions of culture supernatants provided further support to identify candidate metabolites responsible for antimicrobial activity. The *C. albicans* inhibiting vesicles from WAC00276 contained a peak with molecular ions and expected fragmentation pattern from the polyene candicidin (Fig 2A-C)^45^. WAC00288 vesicles, with *S. aureus* inhibitory activity, contained 3 dominant peaks with clear molecular signatures for anthracyclines (Fig 2G-I), namely hexaglycosylated β-rhodomycin analogs from the cosmomycin/ditrisarubicin family, the major one being ditrisarubicin C (Fig 2I, Supplementary Fig 1E,F)^46–48^. The *S. aureus* inhibitory vesicles from the soil isolate WAC00237 had an abundant peak with the molecular ion and expected fragmentation pattern of actinomycin X2 (Fig 2D-F)^49^. The metabolites eluted at very different retention times off the reverse-phase column, highlighting the wide spectrum of polarity and structural diversity in the compounds (Fig 2).

**Figure 2.**
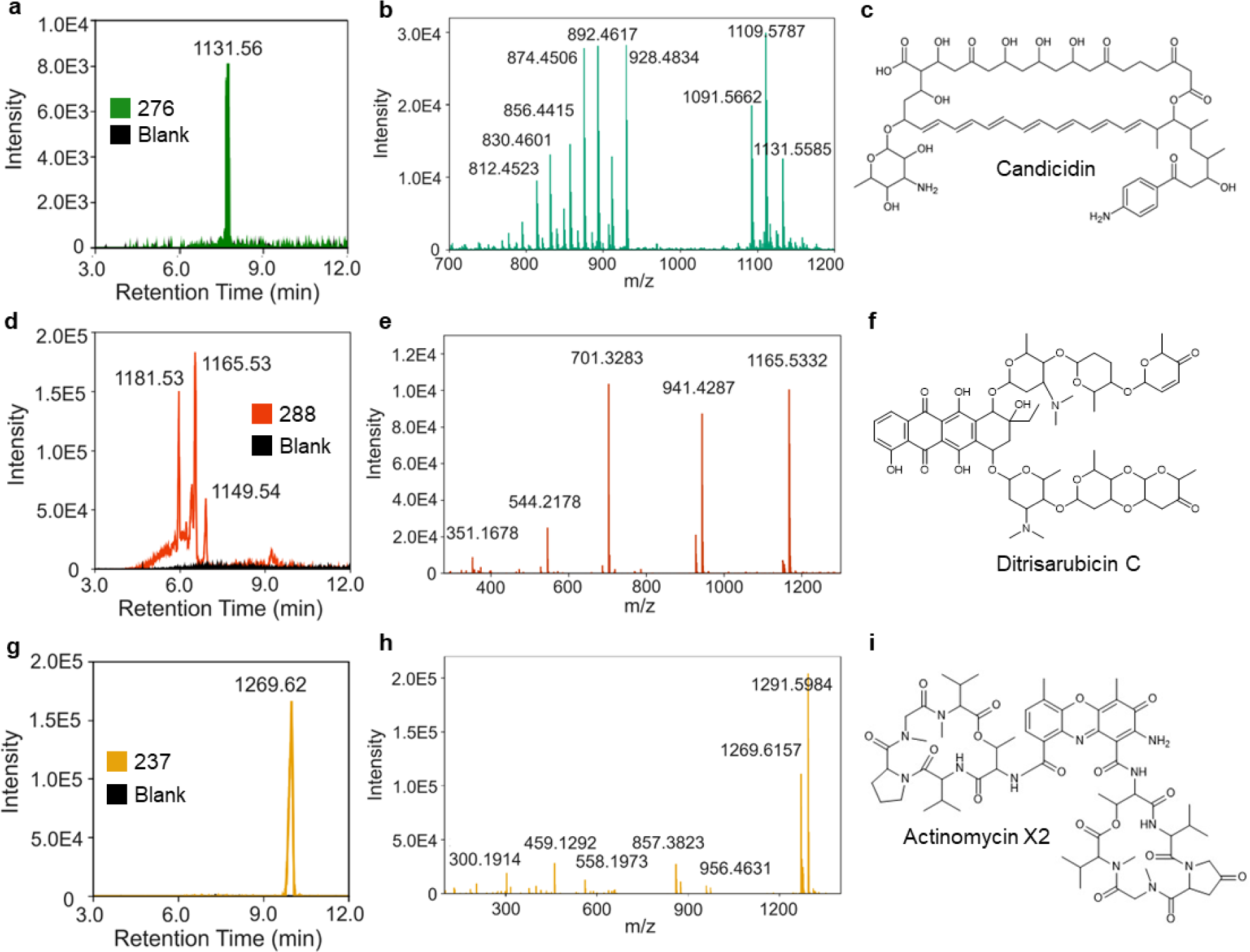
Vesicles package diverse antimicrobial metabolites. LC-MS/MS analysis identified clear peaks present in vesicle extracts (purified by size exclusion chromatography) not present in blanks (black). In WAC276 vesicles (green) a peak was present that matched the expected m/z for the sodium adduct of candicidin (M+Na 1131.56, (A) extracted ion chromatogram of 1131.561 +/− 0.005), and had the expected mass fragments (B) (MS^e^ spectrum of M+Na 1131.56 peak at rt 7.4 min in HPLC purified extracts of WAC276 supernatant) of candicidin (C). WAC288 vesicles (orange) contained several peaks arising from related molecular structures, with m/z of M+H ions of 1181.53, 1165.53, and 1149.54 (D, extracted ion chromatogram of 1100 to 1200), and the expected mass fragments of hexaglycosylated β-rhodomycins (E, MS^e^ spectrum of peak at rt 6.5 min), including ditrisarubcin C (F). WAC237 vesicles (yellow) contained a peak matching the m/z of the molecular ion of actinomycin X2 (M+H 1269.62, (G) extracted ion chromatogram of 1269.621 +/− 0.005), with fragments (H, MS^e^ spectrum of peak at rt 10 min) matching those expected from the molecular structure (I). Results are representative of at minimum biological duplicate batches of vesicles.

These vesicle-packaged antimicrobials differ in chemical structure, bioactivity, and biosynthetic pathway (Fig 1,2, Supplementary Fig 1). In agreement with previous reports, but under different culture conditions, our results confirm that *Streptomyces coelicolor* packages actinorhodin into vesicles^18,43^. Actinorhodin is a MW 634.5 Da, redox-active, planar molecule produced by polyketide synthases and has antimicrobial activity against some Gram-positive bacteria, likely targeting the lipid II pathway^50,51^. Our finding that soil isolate WAC00276 packages the antifungal compound candicidin into vesicles, aligns with the report that the lab-adapted species *S. albus* releases vesicles containing candicidin^22^. Candicidin is produced by many streptomycetes^52^, a MW 1,109 Da amphiphilic polyketide that interferes with ergosterol in fungal membranes. We further identify that soil isolate WAC00237 packages actinomycin X2 MW 1,269 Da into vesicles. Actinomycins are large hydrophobic non-ribosomal peptides containing a core phenoxazinone and two peptide lactone rings. This core can intercalate into DNA leading to inhibition of RNA polymerase. Finally, we find that soil isolate WAC00288 vesicles package several hexaglycosylated anthracyclines (cosmomycins/ditrisarubicins) into vesicles. Anthracyclines are also DNA-intercalating, but chemically distinct to actinomycins. These hydrophilic, highly glycosylated anthracyclines are assembled by polyketide synthases. The diversity of these identified extracellular vesicle packaged compounds, combined with the reports of lipophilic undecylprodigiosin and linearmycins in vesicles from streptomycetes^19,20,43^, suggests that vesicle packaging is permissive in nature, and can be used to package hydrophilic, amphiphilic, and hydrophobic compounds from diverse synthetic pathways, regardless of each compound’s ultimate molecular target or mode of biosynthesis.

In answer to our first question, we conclude that extracellular vesicle packaging of antimicrobial specialized metabolites is widespread in filamentous actinomycetes. This occurred in approximately 50% of strains we cultured, nine out of seventeen strains in our study packaged a significant portion of secreted antimicrobials into vesicles (Fig 1A, Supplementary Table 1). This is remarkable because we used a single screening growth medium and time-point and made no effort to optimize the expression of metabolites or vesicles. Specialized metabolite yields vary considerably with growth condition in the laboratory, and many are not expressed at all^12,53–55^. Given the substantial association with vesicles we observed, we suspect vesicle packaging of specialized metabolites is very common and possibly a ubiquitous capability in the actinomycetes. This is a dramatic departure from a decades-old paradigm of specialized metabolites acting primarily as secreted molecules in solution.

### Specialized metabolite packaging is coincident with and responsible for extracellular vesicle antimicrobial activity

We next examined the temporal association between the secretion of antimicrobial metabolites and of vesicles. Candicidin, anthracyclines, and actinomycins each have notable UV-vis absorbance properties with strong absorption for candicidin at 390 nm, the anthracyclines at 495 nm, and actinomycin X2 at 445 nm, enabling spectrophotometric detection. Culture supernatants of environmental isolates WAC00276, 00288, and 00237 were sampled over time. WAC00276 and WAC00288 high MW fractions had detectable lipids, specialized metabolites, and antimicrobial activity appear on day 2 and continue to climb over the week time-course (Fig 3A,B). In contrast, WAC00237 high MW fractions did not contain significant lipids or metabolites until day 4 (Fig 3A,B). For all strains, the levels of lipids were notably coincident and correlated with the levels of metabolites in the production phase (Fig 3C); antimicrobial activity also tracked closely with the levels of metabolites (Fig 3D). The fact that lipids appeared and increased concomitantly with metabolites in the three different strains, but at different times and rates between the strains, suggests a close link between vesicle production and specialized metabolite production. From the high correlation between antimicrobial activity of the vesicles and the levels of metabolites, we also infer that the antimicrobial activity is primarily attributed to the antimicrobial metabolites packaged.

**Figure 3.**
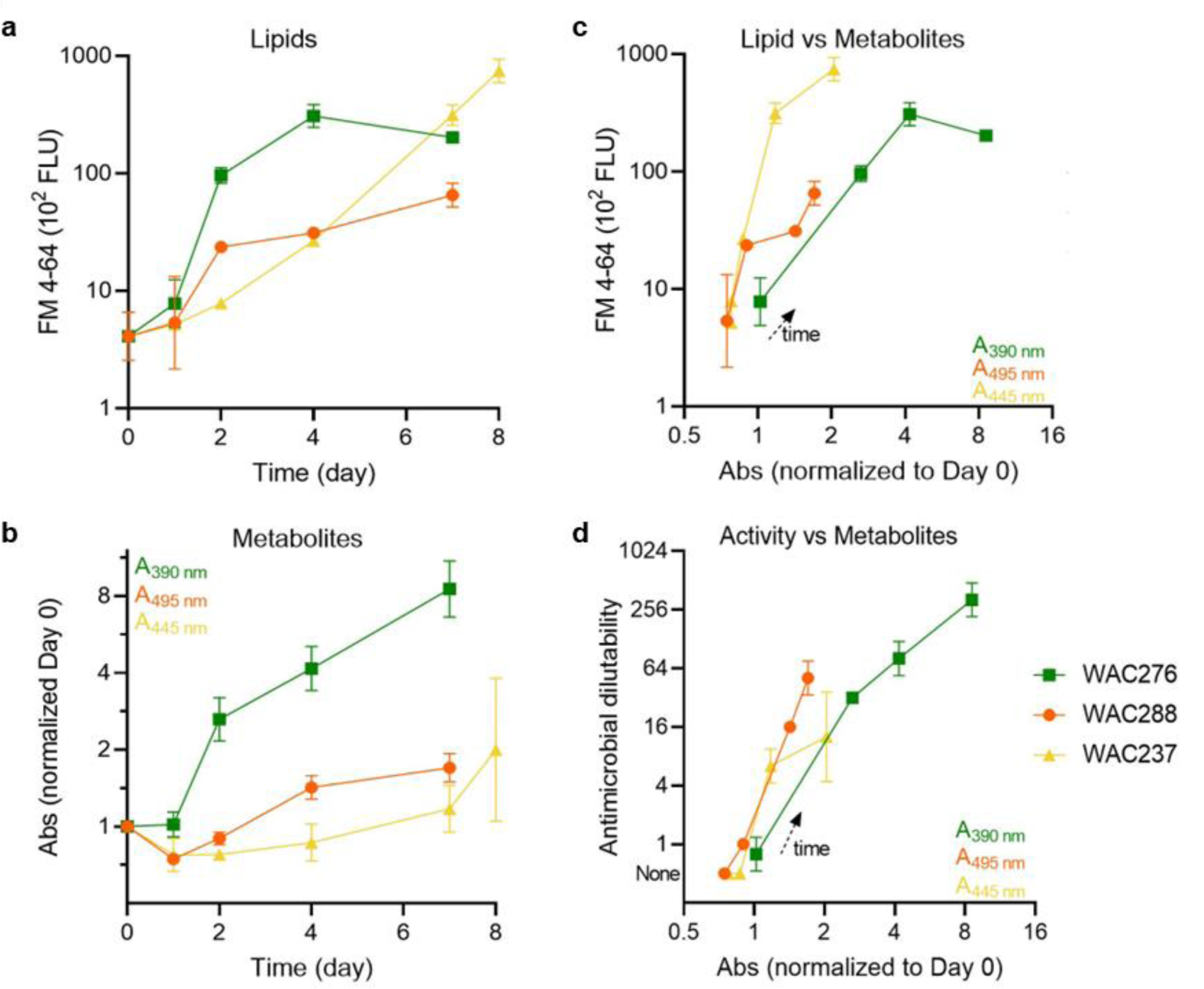
Lipids, metabolites, and antimicrobial activity are coincident in the high molecular weight secretions. WAC276 (green squares), WAC288 (orange circles), and WAC237 (yellow triangles) culture supernatants were sampled over time, the high molecular weight (>100 kDa) fraction purified, and tested for presence of lipids, antimicrobial metabolites, and antimicrobial activity. (A) Lipid presence was indicated by fluorescence of the lipid reactive dye FM 4-64, and (B) metabolites were measured via the absorbance at 390 nm for candicidin (in WAC276), 495 nm for anthracyclines (in WAC288), and 445 nm for actinomycin (in WAC237) in butanol extractions of the high molecular weight fractions. (C) Lipid signal increased coincidently with the metabolite level, (D) as did the antimicrobial activity (WAC276 fractions against *C. albicans*, WAC288 and WAC237 against *S. aureus*, inhibition of growth of liquid cultures of serial dilutions). Results are technical triplicate from an experiment representative of time-courses done in biological triplicate.

To answer the question of whether vesicle production and activity requires the presence of specialized metabolites, we focused on the anthracyclines from soil isolate WAC00288. These hexaglycosylated β-rhodomycins have been described as antibacterial, antifungal, antiphage and insecticidal, covering a broad range of predators filamentous actinomycetes may encounter in the environment, and are hence of considerable interest^46,47,56,57^. They are composed of a core anthraquinone decorated with six sugar chains (Fig 1D), and their antimicrobial and cytotoxic activity depends on their ability to intercalate DNA^48,56^. We took advantage of modified strains of WAC00288 in which key regulatory and biosynthetic genes required for the anthracycline (cosmomycins/ditrisarubicins) production are inactivated^47^. These knockout strains were cultured in parallel with the wild type parent. We confirmed the absence of anthracyclines in all three mutant supernatants by mass spectrometry. Vesicles were still produced, determined by lipid signal, nanoparticle tracking analysis, transmission electron microscopy, and SEC purification (Fig 4A, B, Supplementary Fig 2). Interestingly there was a 10-fold reduction in final vesicle numbers in the mutant strains relative to the wildtype (Fig 4A, Supplementary Fig 2). There was also a complete loss of antimicrobial activity in both high and low molecular weight fractions (Fig 4C). This loss of antimicrobial activity in the mutant vesicles, but clear connection between the amount of anthracyclines and antimicrobial activity in the wildtype vesicles (Fig 3), demonstrates that the anthracyclines are both required and responsible for vesicle antimicrobial activity. Moreover, when wildtype WAC00288 high molecular weight fractions were treated with detergent to lyse the vesicles, anthracyclines and antimicrobial activity were then recovered from the low molecular weight fraction (Supplementary Fig 3).

**Figure 4.**
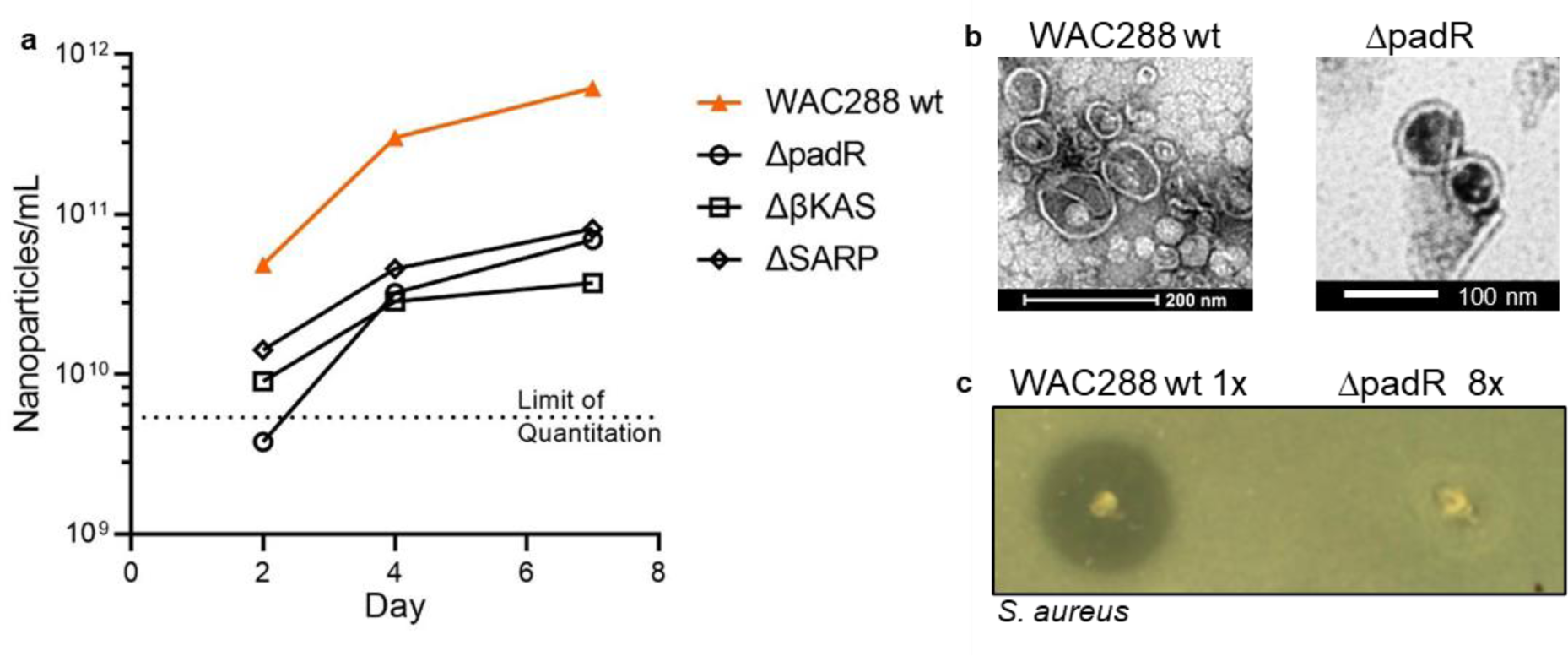
Anthracycline production increases vesicle yields and is required for antimicrobial activity of WAC288 vesicles. (A) High molecular weight fractions purified over time from wildtype (wt, orange triangles) and three WAC00288 strains knocked out for different essential genes in anthracycline (cosmomycin/ditrisarubicin) production, a padR regulator, a βKAS synthetic gene, and a SARP regulator, (black open symbols), were assayed for nanoparticle concentration per mL by nanoparticle tracking analysis. (B) Vesicles purified from wt and the padR knockout by SEC were imaged by negative stain transmission electron microscopy. (C) WAC288 wt high molecular weight fractions create zones of inhibition against *S. aureus*, but fractions from the padR knockout do not, despite concentration of 8 fold to match the vesicle numbers of the wt. Results are representative of biological duplicate.

In conclusion the production of metabolites and of extracellular vesicles was highly associated in our strains, following different temporal profiles between isolates (Fig 3, 4). Through rigorous purification and genetic knock-outs, we demonstrated that the anthracycline cargo was essential for the antimicrobial activity of WAC00288 extracellular vesicles (Fig 4); however, the metabolites were not required for extracellular vesicle production. Interestingly, the total yield of vesicles was higher when the cargo was present. This is similar to previous observations in *Streptomyces sp. Mg1* and *Pseudomonas aeruginosa* where strains bearing deletions of the linearmycin or PQS cargo genes had reduced vesicle yields during purification^20,37^. In contrast, a violacein null mutant of *Chromobacterium violaceum*^58^ produced greater vesicle yields relative to the wild type. It was argued that the hydrophobicity of these cargo might alter vesicle production by modulating cellular membrane dynamics^20,37^. This seems less likely for the hydrophilic and highly glycosylated anthracyclines. An alternative view would be that in some cases, rather than influence the secretion of vesicles, the metabolite cargo enhances vesicle stability and recovery on purification. Given the wide variety of chemical cargo in extracellular vesicles that we (Fig 2) and others have observed, and this repeated observation of a close association of vesicle production with production of specialized metabolite cargo (Fig 3, 4), there may be regulatory mechanisms that link metabolite and vesicle biogenesis pathways.

### Extracellular vesicles deliver antimicrobials to cells via membrane fusion

We next turned to examine the effect vesicle encapsulation has on the transport and delivery of specialized metabolites. We hypothesized that extracellular vesicles might deliver their antimicrobial metabolite content to target cells by direct fusion with cellular lipid membranes. This is particularly important for specialized metabolites with a cytoplasmic location of action, such as the anthracyclines which intercalate into the DNA. The transfer of lipid soluble fluorescent dyes is one method used to assess extracellular vesicle fusion with membranes^59^. Labeled and unlabeled vesicles (SEC-purified) were incubated with target cells. The cells were then pelleted and washed at low centrifugal forces that do not pellet vesicles. Two methods were used to compare fluorescence of treated cells to negative controls of cells or vesicles alone, confocal microscopy and spectrometry (Fig 5, 6).

**Figure 5.**
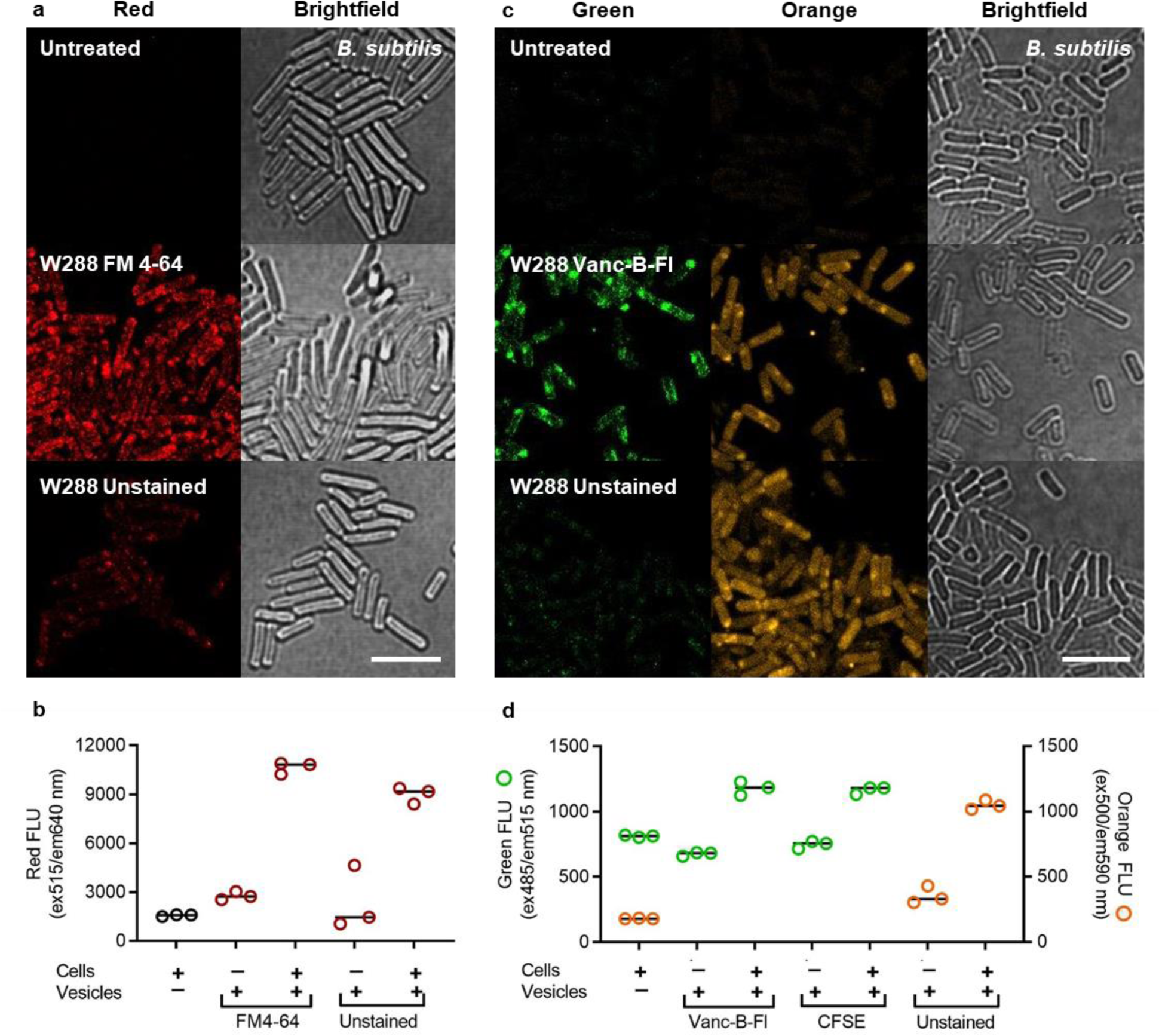
WAC00288 vesicles deliver multiple fluorescent metabolites to membranes or cytoplasm of *B. subtilis*. *B. subtilis* cells were incubated for 1 h with vesicles from WAC288 (W288), either labeled with external fluorescent dyes or unstained with native cargo of fluorescent anthracyclines. Cells were pelleted (5000xg, 5 min) and washed 100 fold with saline, then assessed for fluorescence. For microscopy, cells were placed on an agarose pad and imaged by confocal microscopy (Red ex488 nm, em650 – 760 nm; Green ex488 nm, em490 – 550 nm; or Orange ex488 nm, em570 – 650 nm; and brightfield, T-PMT) (A, C). For spectrophotometer readings, cells were transferred to microtiter plate wells (Red ex515 nm, em640 nm; Green ex485 nm, em515 nm; Orange ex500 nm, em590 nm) (B, D). Images grouped in panels are from parallel treatments in one experiment, placed on one agarose pad, imaged and processed with the same settings. White bar, 5 µm. *B. subtilis* cells incubated for 1 hr with unstained WAC00288 vesicles or equal amounts of FM 4-64 (A, B), Vancomycin-Bodipy-Fl (Vanc-B-FL) (C, D), or CFSE (D) labeled vesicles, acquired fluorescence corresponding to the various molecular cargo. (B,D) Results shown are from technical triplicate of an experiment representative of experiments done in at least biological duplicate.

Vesicles from WAC00288 carrying the lipid soluble dye FM 4-64 were able to transfer the red fluorescence to *B. subtilis* cells (Fig 5A, B). This transfer of fluorescence was significantly greater than the negative controls of cells alone, stained vesicles alone, or an equal amount of unstained vesicles incubated with *B. subtilis* (Fig 5A, B). Fluorescence was transferred to stationary or actively growing cells and could be detected even at the shortest incubation time tested of 15 min (Supplementary Fig 4). Microscopy revealed the FM 4-64 fluorescence in a cell membrane associated pattern on the *B. subtilis* (Fig 5A). To validate this with a second molecule of different chemical composition^60^, WAC00288 vesicles were labeled with vancomycin conjugated to bodipy-fluorescein; this binds tightly to lipid II in bacterial membranes through the vancomycin moiety. Again, WAC00288 vesicles were able to transfer the green vancomycin fluorescence to *B. subtilis,* and in a membrane associated pattern (Fig 5C, D). This vesicle-mediated transfer and delivery of two distinct lipid-associated dyes to cells, strongly supports direct vesicle fusion to the cell membrane.

We then asked if the vesicle fusion could deliver the native cargo of specialized metabolites. Anthracyclines have intrinsic fluorescent properties. We reasoned that the hexaglycosylated β-rhodomycins are likely to be carried in the vesicle lumen due to the hydrophilicity of the six sugar moieties and the aqueous solubility of the purified compounds (Fig 2B). Unstained, anthracycline-containing WAC00288 vesicles were incubated with *B. subtilis*. Notably we found that the *B. subtilis* cells acquired anthracycline fluorescent properties (Fig 5C, 5D). Confocal microscopy of the cells revealed anthracycline associated fluorescence (orange), which was diffuse throughout cells suggesting cytoplasmic localization (Fig 5C). This cytoplasmic location was especially clear in vancomycin stained vesicle transfer experiments, where the native anthracycline orange fluorescence could be imaged in parallel to the green vancomycin fluorescence (Fig 5C), and in comparison to fluorescence transfer from anthracycline-knockout WAC00288 vesicles (Supplementary Fig 5). As a further test of the ability to transfer aqueous lumenal content, WAC00288 vesicles were incubated with the membrane permeable carboxyfluorescein diacetate succinimidyl ester, which is converted by cytoplasmically localized esterases to amine-bound fluorescent CFSE. Vesicles became fluorescent with CFSE and were able to transfer green CFSE fluorescence to *B. subtilis* cells (Fig 5D).

Altogether this transfer of two distinct lipid associated molecules, and two distinct aqueous molecules, confirms extracellular vesicle fusion to cell membranes and delivery of molecular cargo. Importantly, these results also demonstrate direct delivery of vesicle cargo anthracyclines into target microbial cell cytoplasm, the cellular location necessary for antimicrobial activity.

### Vesicles provide a mechanism to transfer antimicrobials to diverse microbial life forms

To determine if vesicle fusion with membranes could mediate antimicrobial delivery to a variety of organisms, including problematic pathogens, we incubated the streptomyces vesicles with further microbes, including the Gram-positive bacteria, *S. aureus*, the Gram-negative bacteria *K. pneumoniae*, and the yeast *Cryptococcus neoformans*. Strikingly WAC00288 vesicles, either labeled with FM 4-64 or unstained carrying fluorescent anthracyclines, were able to transfer both the lipid associated FM 4-64 fluorescence, and anthracycline associated fluorescence to all microbes (Fig 6A-F, Supplementary Fig 6). Similarly, we purified WAC00237 or WAC00276 vesicles labeled with FM 4-64 and incubated them with *S. aureus* or *C. albicans* respectively. Again, we found that the *S. aureus* and *C. albicans* acquired membrane associated FM 4-64 (Fig 6G-J). In *C. neoformans* and *C. albicans* the FM 4-64 is seen labeling the endocytic vesicles of the yeast due to the high endocytic turnover of yeast cellular membranes^61^.

**Figure 6.**
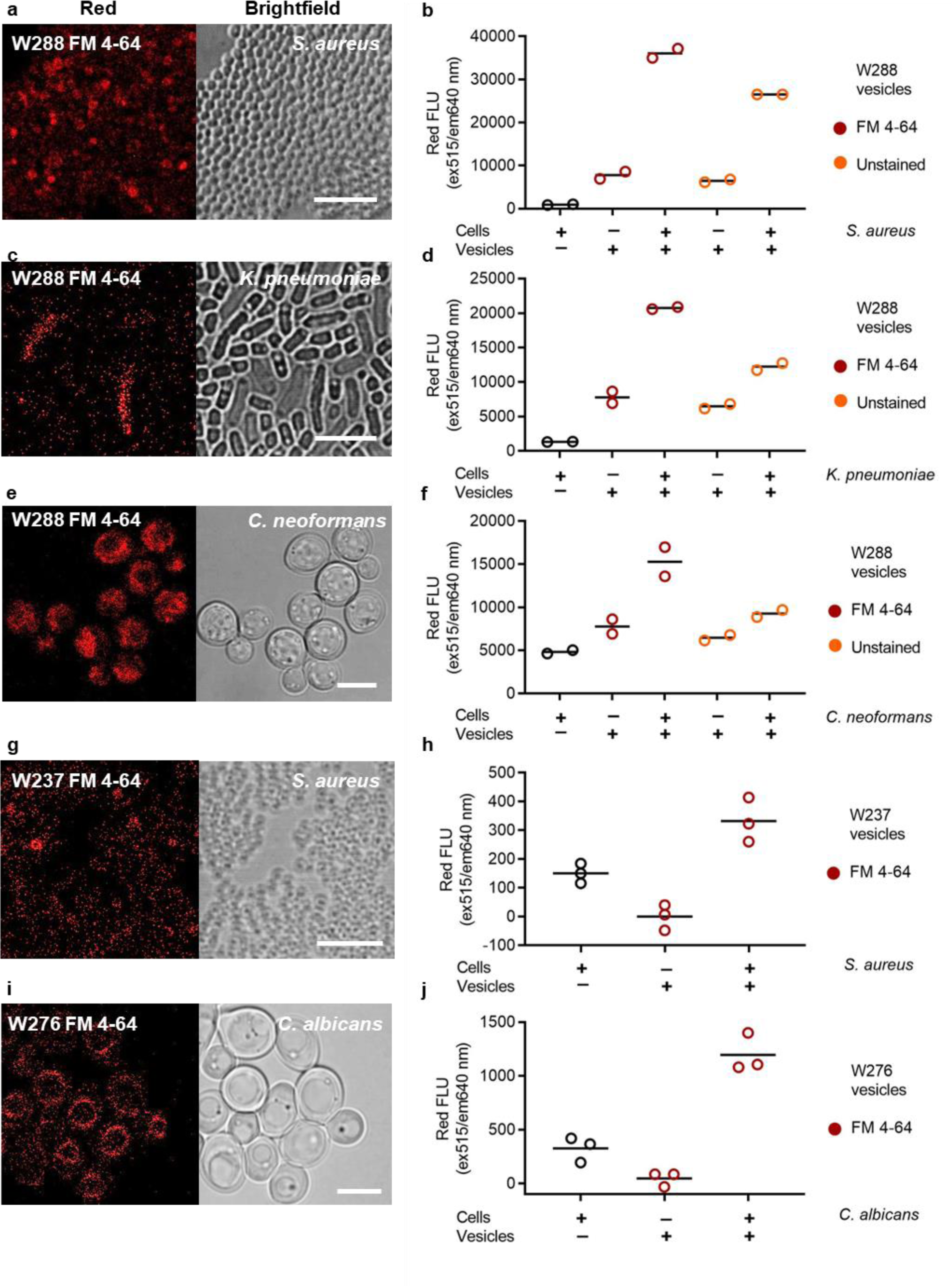
Streptomyces vesicles are a broad spectrum delivery mechanism for molecular cargo to pathogenic bacteria and yeast. Labeled (FM 4-64, red circles) or unstained (orange circles) vesicles from WAC288 (A – F), WAC237 (G, H), or WAC276 (I, J) were incubated with microbial cells, *S. aureus* (A, B, G, H), *K. pneumoniae* (C, D), *C. neoformans* (E, F), or *C. albicans* (I, J), and were able to transfer fluorescence from the molecular cargo to each cell type. Cells were pelleted (5000xg, 5 min) and washed 100 fold with saline, then imaged by confocal microscopy (Red, ex488 or 515 nm, em 650 - 760 nm), and assayed for fluorescence with a spectrophotometer (Red ex515 nm, ex640 nm). Controls for expected background fluorescence were vesicles or cells alone. Results shown are technical replicates from an experiment representative of at least biological duplicate.

We conclude *Streptomyces* extracellular vesicles are able to transfer fluorescence from multiple lipid and aqueous soluble molecules to diverse prokaryotic and eukaryotic microbes, most importantly from naturally packaged antimicrobials. This is strong support for the hypothesis that extracellular vesicles can serve as a delivery mechanism for specialized metabolite cargo via membrane fusion to other microbial cells. Notably, we find a versatile, broad and permissive spectrum for fusion and content delivery. We demonstrated the transfer of multiple molecules (lipid carried vancomycin and FM 4-64, and lumenal anthracyclines and CFSE) from *Streptomyces* vesicles to other microbes. Metabolites can go to either the membrane (vancomycin) or the cytoplasm (anthracyclines) (Fig 5), enabling interaction with molecular targets. This vesicle fusion and delivery was seen from three different environmental isolates, WAC00276, 00237, and 00288 (Fig 6), suggesting it is a general property of *Streptomyces* extracellular vesicles. Remarkably membrane fusion is able to occur in Gram-positive and Gram-negative bacteria as well as fungi (Fig 6). Recent advances have highlighted the porous and heterogeneous nature of microbial cell envelopes, along with rapid shifts in membrane dynamics, presenting possibilities for how extracellular vesicles can both be released from and fuse with cell membranes^62^.

## Conclusion

In summary, we examined extracellular vesicle packaging of antimicrobial specialized metabolites in filamentous actinomycetes, and found that antimicrobial vesicle production is widespread, occurring in over 50% of strains we cultured (Fig 1). Critically, we found that purified vesicles from different strains are associated with diverse specialized metabolites, and have various spectrums of antimicrobial activity against bacteria and fungi (Fig 2). We found that vesicles form independently of the antimicrobial metabolites themselves, though the presence of the metabolite cargo is coincident with vesicle production and can increase vesicle yield (Fig 3, 4). We demonstrate that vesicles fuse with prokaryotic and eukaryotic microbial cells, delivering small-molecule cargo to either the cell membrane (FM 4-64 and vancomycin) or the cytoplasm (anthracyclines and CFSE) (Fig 5, 6). Crucially, fusion was achieved in *S. aureus, K. pneumoniae*, *C. neoformans*, and *C. albicans*, each of which are priority pathogens that cause infections highly resistant to available antibiotics. These striking features have implications for how we understand the action of specialized metabolites in nature and for how to use them clinically.

This direct packaging and delivery of a broad variety of metabolites to a broad variety of microbes has important implications for the way we understand the action of *Streptomyces* specialized metabolites in the environment. Extracellular vesicle packaging and release of specialized metabolites would provide protection of the metabolites from extracellular degradation, and fusion to other cells would impact the dynamics of bioactivity. Vesicle packaging can also inform translational development. The vast majority of specialized metabolites that have been developed for clinical use have been developed as orally bioavailable drugs or injectable solutions, with the desired outcome being systemic distribution of soluble compound. The direct delivery via vesicle fusion of diverse antimicrobial specialized metabolites that we observed, particularly to notoriously antimicrobial resistance species such as *Klebsiella, Cryptococcus* and *S. aureus* (Fig 6), suggests future therapeutic avenues. This lends credence to the development of either extracellular vesicles themselves^63–66^, or lipid nanoparticle formulations^67,68^, as therapeutic vehicles for the delivery of small molecule antimicrobials to their targets. Indeed, the clinically used polyene amphotericin, closely related to candicidin (Fig 2A), works better in a liposomal formulation; this is mostly attributed to improved pharmacokinetic and toxicity properties^69^; however it has also been suggested that the liposomal formulation improves delivery of the drug to the fungal membranes^70^. With the benefits, safety, and feasibility of lipid particle formulation most conclusively demonstrated to date with the mRNA Covid-19 vaccines, the time is ripe for lipid formulations of antimicrobials to counter antimicrobial resistant pathogens. We suggest extracellular vesicle packaged specialized metabolites from actinomycetes offer an attractive source of candidate molecules.

## METHODS

All experiments were done at minimum in biological duplicate, with results shown from a representative experiment.

### Actinomycete culture and supernatant size separation

Actinomycete strains were stored as glycerol (18%) spore stocks, environmental isolates were from the Wright Actinomycete Collection^42^. Strains were cultured with Maltose-Yeast Extract-Malt Extract (MYM)^71^ medium, on agar or in liquid. For the antimicrobial vesicle screen, fresh spores were collected from one week agar cultures, and 10 mL of liquid MYM inoculated with approximately 5×10^6^ CFU/mL then cultured for one week. WAC00237, WAC00276, and WAC00288 wild-type and knock-out strains^47^ were stored as aliquoted glycerol spore stocks and liquid MYM cultures inoculated directly from aliquots with 5×10^6^ CFU/mL. After incubation at 30°C with shaking at 220 rpm, cells were removed by centrifugation (3000xg, 20 min) and filtration (0.2 μm). Supernatant was either used fresh or stored at −20°C, then separated into high molecular weight and low molecular weight fractions with 100 kDa 0.5 mL centrifugal filters (Amicon Ultra) at 9500xg, concentrating the high molecular weight fraction 10-fold before washing 100-fold with saline. Low molecular weight filtrates were dried on a Genevac (EZ-2) for 1 h, before resolubilizing in water for 10-fold concentration.

### Lipid assays with FM 4-64

Fluorescence was measured at excitation 515 nm, emission 640 nm (Synergy H1, Biotek). Matched high and low molecular weight fractions adjusted to supernatant concentration equivalents in 0.9% saline were measured for fluorescence before and after addition (15 min incubation) of 5 μg/mL FM 4-64 (Invitrogen), and the background reading subtracted to obtain the final measurement.

### Test microbial culture and antimicrobial assays

*Staphylococcus aureus* ATCC 25912 (37°C), *Klebsiella pneumoniae* ATCC 13883 (37°C), and *Bacillus subtilis* 168 (30°C) were grown with Mueller Hinton II medium; and *Candida albicans* ATCC 90028 and *Cryptococcus neoformans* H99 were grown with Yeast Extract-Peptone-Dextrose medium at 30°C. The initial screen conditions were 10-fold concentrated size fractions from supernatant, 2 µL on lawns, and at 10% v/v in liquid cultures in microtiter plates. Lawns were created from 1:100 dilutions of overnight cultures, liquid cultures were 1:1000 dilutions of overnight cultures, and inhibition was measured after 18 h. To test purified vesicles or soluble fractions (fresh or stored at −80°C), samples were diluted in saline and spotted against lawns, or mixed 1:1 with liquid cultures in at least technical duplicate.

### Transmission Electron Microscopy

Negatively charged carbon grids (CF400CU50, EMS) were incubated with samples for 60 s, washed three times with water, then stained with 2% uranyl acetate for 30 s. Grids were imaged using a Talos L120C transmission electron microscope (software TEM Imaging & Analysis v5.0 SP4).

### Vesicle purification by Size Exclusion Chromatography

Culture supernatant was centrifuged to pellet cells (3000xg, 20 min), filtered through 0.2 μm membranes (PVDF or PES), and used directly or concentrated 5 to 30 fold using a 10 kDa centrifugal filter (Amicon). This concentrated fraction was then split into unstained controls and into portions incubated with fluorescent dyes. FM 4-64 was at 10 μg/mL for 15 min, Vancomycin-BodipyFL at 1 μg/mL for 30 min, and CFSE at 10 μM for 30 min. Samples were then applied in parallel to size exclusion columns (qEV1, 35 nm IZON Sciences) and separated with PBS elution, collecting 1 mL fractions. Fraction 1 to 3 after the void volume (4.5 mL) were combined as the vesicle fraction – these fractions had strong lipid signal from FM 4-64 fluorescence and high particle counts by nanoparticle tracking analysis.

### Nanoparticle Tracking Analysis

Samples were iteratively diluted in saline and analyzed by nanoparticle tracking analysis (NanoSight NS300, Malvern) until particles were at quantifiable concentrations. For time-course comparisons, diluted samples were stored at −20°C then analyzed in one batch to keep acquisition and analysis settings consistent. Three thirty second runs were collected for each sample.

### Metabolite assays and mass spectrometry

For mass spectrometry, 20 μl of size exclusion purified vesicles were incubated with 60 μl of 1:1 isopropanol and acetonitrile for 10 min, then precipitate pelleted by centrifugation at 16,000xg for 10 min, and 10 μl of the supernatant injected on an Acquity UPLC BEH C18 column (1.7 μm, 2.1 × 50 mm). Separation gradient was at 0.2 mL/min, 5% B for 2 min, increasing linearly to 95% over 10 min, and held at 95% for 5 min: A – water + 0.1% formic acid, B – acetonitrile + 0.1% formic acid. Samples were analyzed by electrospray ionization with MS^e^ on a QTOF (Waters Xevo G2S-QTOF). To verify metabolite identity and spectrophotometric characteristics, butanol extracts of supernatants were dried (EZ-2 Elite, Genevac) then resolubilized in 3:1 acetonitrile to water and injected on a C18 analytical column (250x 4.6 mm, 5 µm, 100 Angstrom, Luna, Phenomenex) and separated on an Alliance HPLC (Waters) at 0.8 mL/min using a step gradient of increasing percentage B, from 5% to 90% over 5 min, then holding for a further 7 min: A - water + 0.1% formic acid, B – acetonitrile + 0.1% formic acid. Fractions were dried (Genevac) and resolubilized in DMSO or in water, and the bioactive fractions re-examined by LC-MS/MS, and then assayed for absorbance and fluorescence spectral scans (Synergy H1, Biotek) to determine wavelength maxima. Absorbance maxima for candicidin was 390 nm, anthracyclines 495 nm, and actinomycin X2 445 nm. Fluorescence maxima for anthracyclines was ex 500 nm, em 590 nm. Following this, supernatant or size exclusion purified vesicles were extracted with butanol (1:2) overnight at 4°C and the butanol fraction transferred to microtiter plates for absorbance measurement (Synergy, Biotek).

### Fluorescence transfer assays

Matched unstained and stained vesicle fractions purified by SEC were incubated 1:1 with liquid cultures (overnight cultures were diluted 10- or 100-fold, incubated for 2 h, then mixed with samples) or blank medium, in technical duplicate or triplicate. Cells were pelleted (5000xg, 5 min), washed twice with saline (100x wash), resuspended and fluorescence read in microtiter plates (Synergy H1, Biotek). Excitation, emission (nm): 515, 540 FM 4-64; 500, 590 anthracyclines; 485, 515 Vanc-BodipyFl and CFSE. Experiments were repeated in at least biological duplicate with different batches of vesicles. Results were consistent with vesicles purified fresh or stored at −20°C or −80°C. After a freeze-thaw, vesicle samples were spun at 5000xg for 5 min and supernatant transferred to a clean tube to ensure large aggregates of vesicles were removed.

### Confocal microscopy

Washed cells from fluorescence transfer assays were spotted onto agarose pads (1.5% in saline) and imaged with a 100x oil objective on a Zeiss LSM 880 Elyra. Images were captured and adjusted for brightness and contrast using Zen (v2.1 SP3). For comparative analyses samples were spotted onto the same agarose pad and imaging and software settings were constant between samples. Biological duplicates were imaged for each condition, and all samples had multiple images captured across the spread of the cells on the agarose pad.

## ACKNOWLEDGEMENTS

We thank Karen Maxwell and Anne van der Meij for helpful discussions and comments on the manuscript. Imaging was performed with the support of the Microscopy Imaging Laboratory in the Temerty Faculty of Medicine, University of Toronto, and nanoparticle tracking analysis in the Hospital for Sick Children’s Structural & Biophysical Core Facility. Illustrations were made with Biorender. Funding was provided by the Canadian Institutes of Health Research, Fellowship MFE-176478 to KJM, grant MID-406688 to JRN.

## AUTHOR CONTRIBUTIONS

Conceptualization, K.J.M and J.R.N; Methodology, and Investigation, K.J.M; Resources, J.R.N; Supervision, J.R.N; Writing – Original Draft, K.J.M; Writing – Review & Editing, K.J.M and J.R.N; Funding Acquisition, K.J.M and J.R.N.

---

**Supplementary Figure 1.**
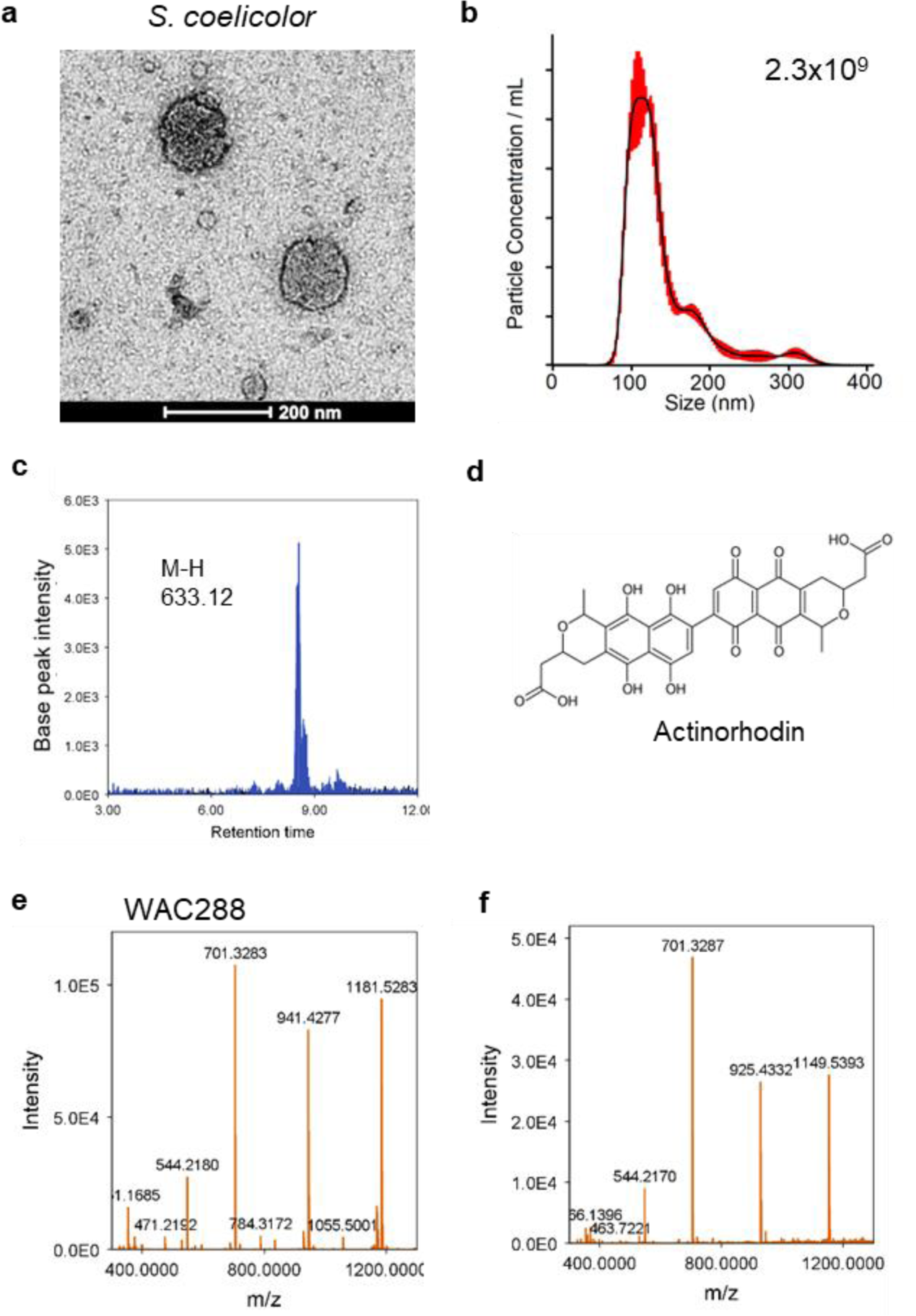
Antimicrobial specialized metabolites are associated with extracellular vesicles from filamentous actinomycetes. (A) *Streptomyces coelicolor* supernatant high molecular weight (MW) fraction >100 kDa was imaged by negative stain transmission electron microscopy revealing extracellular vesicle structures, (B) quantified by nanoparticle tracking analysis, (C) and analyzed by LC-MS/MS (ESI negative mode) determining presence of actinorhodin (structure shown (D) by extracted ion chromatogram of molecular ion m/z (M-H 633.12) in the vesicle extract (blue) but not present in blank (black). (E,F) Hexaglycosylated β-rhodomycin fragments seen in MS^e^ LC-MS/MS spectra at 5.9 min (left) and 6.9 min (right) of WAC00288 vesicles.

**Supplementary Figure 2.**
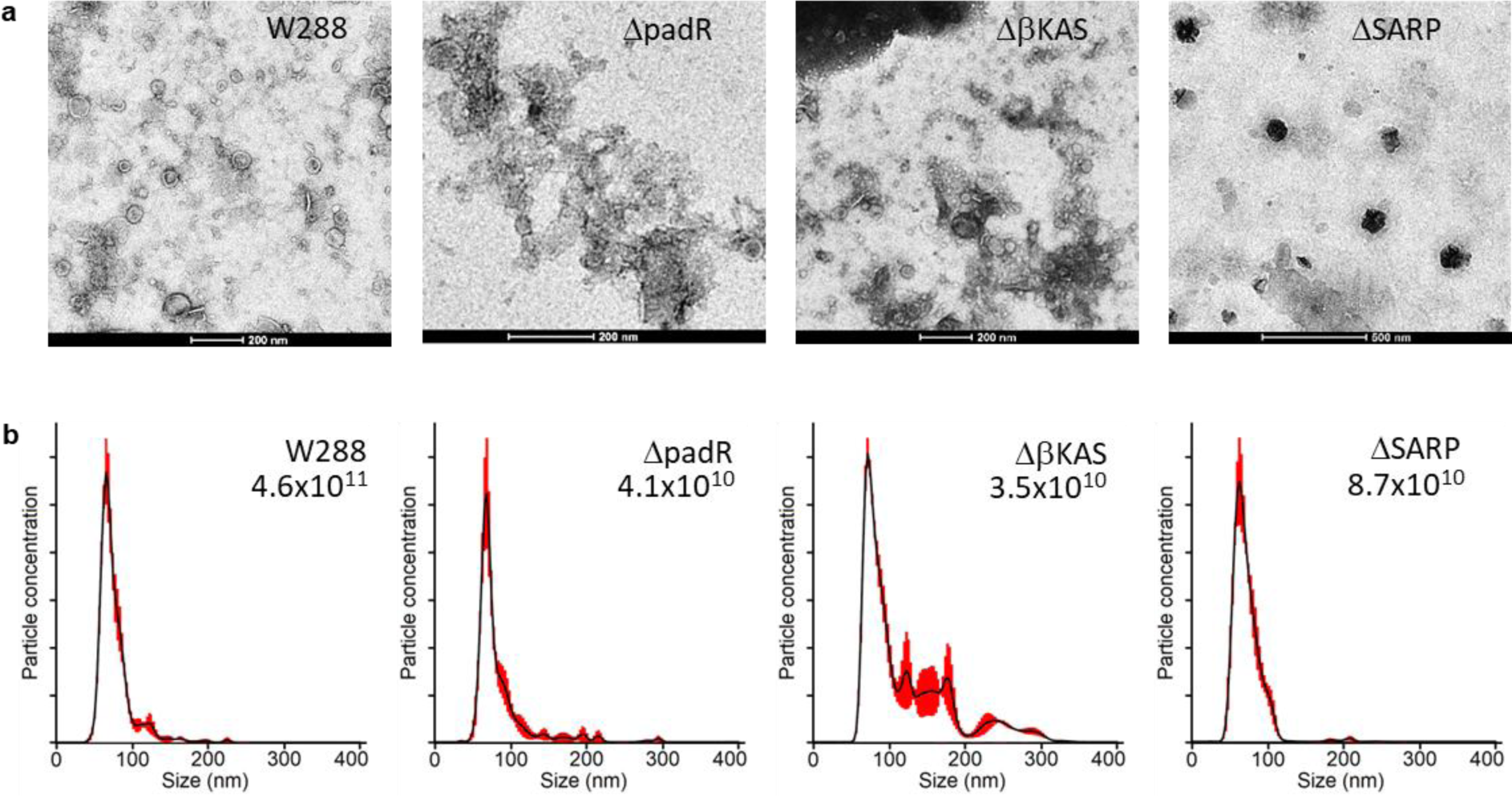
WAC00288 cells knocked out for anthracycline production produce extracellular vesicles, but fewer. WAC00288 (W288) wildtype and strains knocked out for essential genes for anthracycline production – a padR regulator, a βKAS synthesis gene, and a SARP regulator – were cultured in parallel and high MW fractions (>100 kDa) collected and imaged by negative stain transmission electron microscopy (A), or vesicles were purified by size exclusion chromatography and quantified by nanoparticle tracking analysis (B) for particle concentration (displayed per mL relative to unconcentrated supernatant, top right inset), and for size distribution.

**Supplementary Figure 3.**
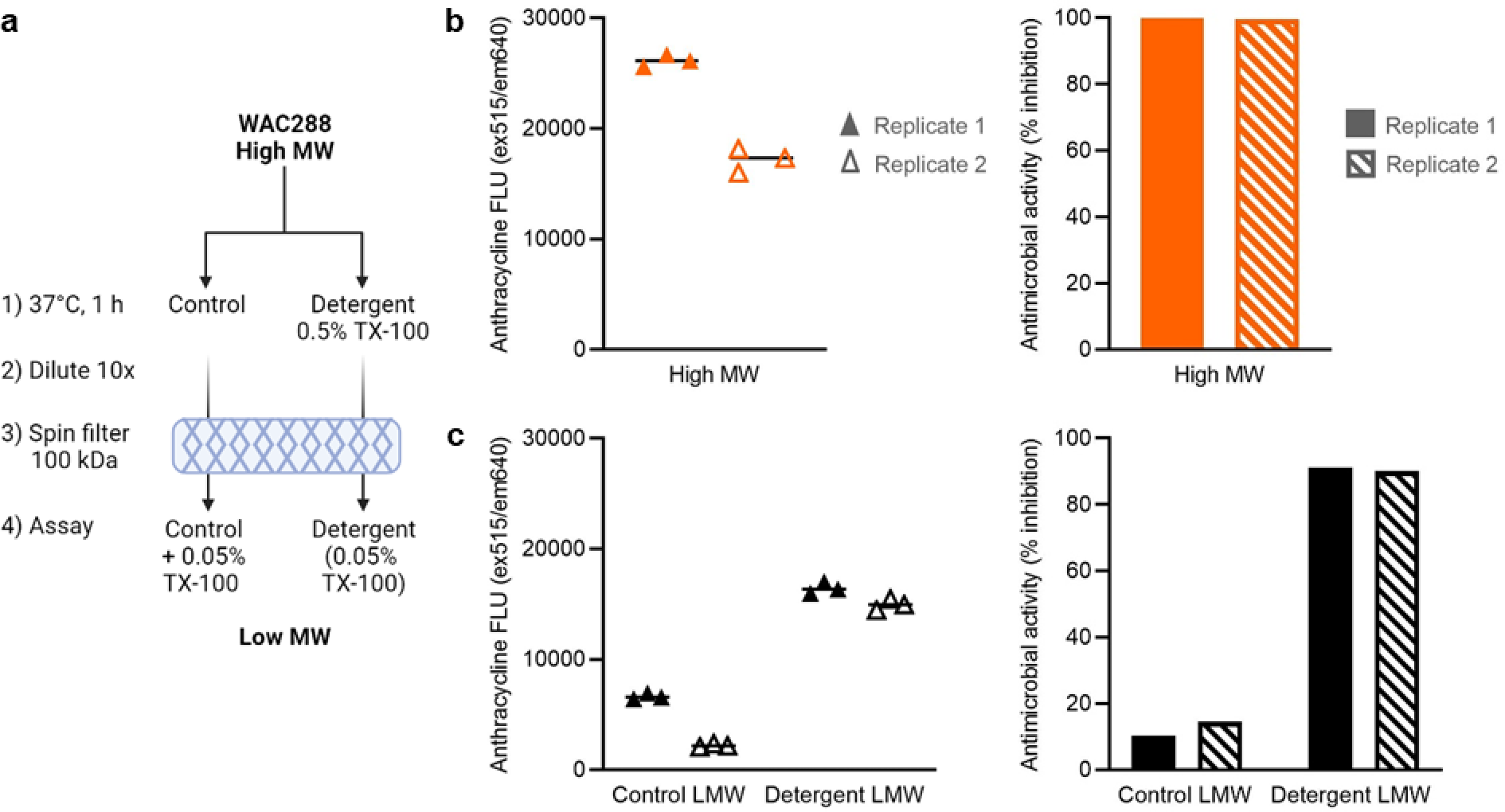
Antimicrobial anthracyclines in extracellular vesicles in WAC00288 high molecular weight fractions. (A) High molecular weight (>100 kDa) fractions from WAC00288 supernatants were partitioned, and 0.5% triton X-100 added to one half, then incubated at 37°C for 1 h. After incubation fractions were diluted 10x in saline to reduce detergent concentration and filtered through a 100 kDa spin column. The low molecular weight (<100 kDa) filtrate of the control had 0.05% triton X-100 added to match the detergent treated filtrate. The anthracycline content (red fluorescence ex515,em640 nm) and antimicrobial activity (percent inhibition of *S. aureus* cultures, OD600) were assayed for the intact high molecular weight fractions (orange, B), and the processed low molecular weight fractions (black, C) in technical triplicate for two biological replicates (filled triangles and bars, and open triangles and bars respectively).

**Supplementary Figure 4.**
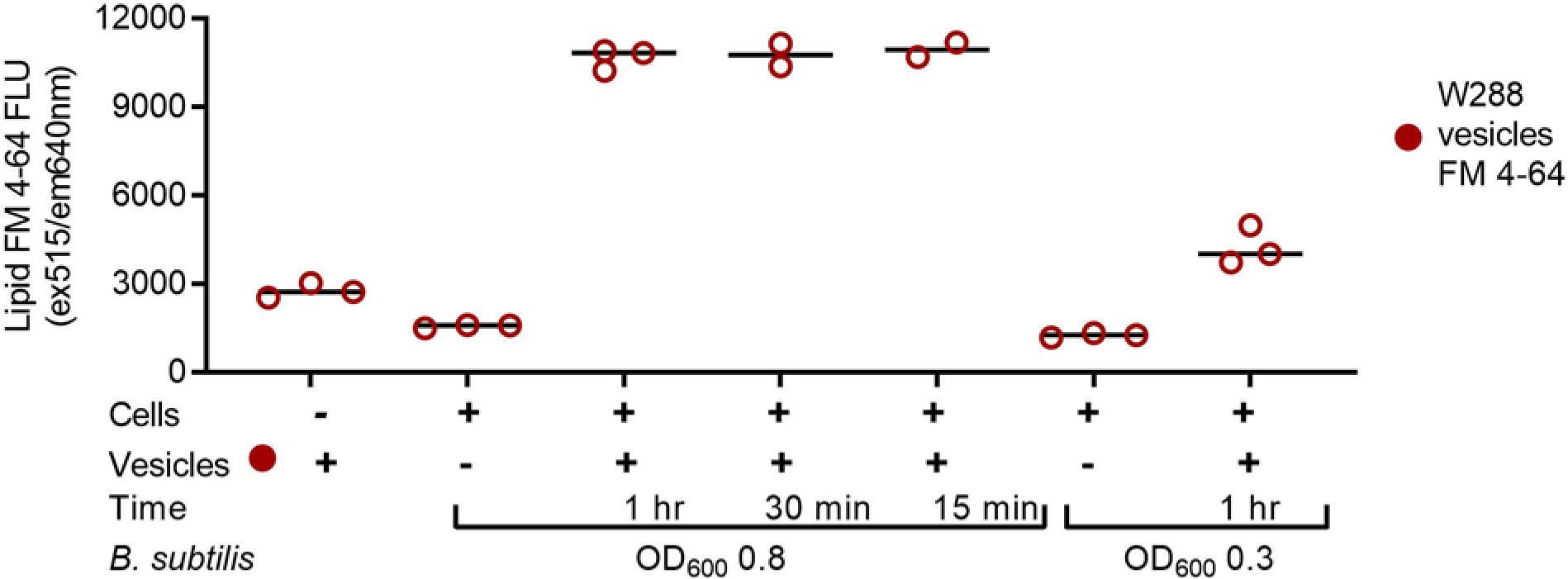
WAC00288 vesicles are able to quickly transfer fluorescence to stationary or exponential Bacillus. FM4-64 labeled WAC288 vesicles (red solid circle) were incubated with microbial cells. Cells were pelleted (5000xg, 5 min) and washed 100 fold with saline, then assayed for fluorescence (open circles). Controls for expected background fluorescence were vesicles or cells alone (open circles). Vesicles were incubated with *B. subtilis* cells at OD_600_ 0.8 for 1 hr, 30 min, or 15 min, or for 1 hr with cells at OD_600_ 0.3, fluorescence measured at ex 515, em 640 nm (red).

**Supplementary Figure 5.**
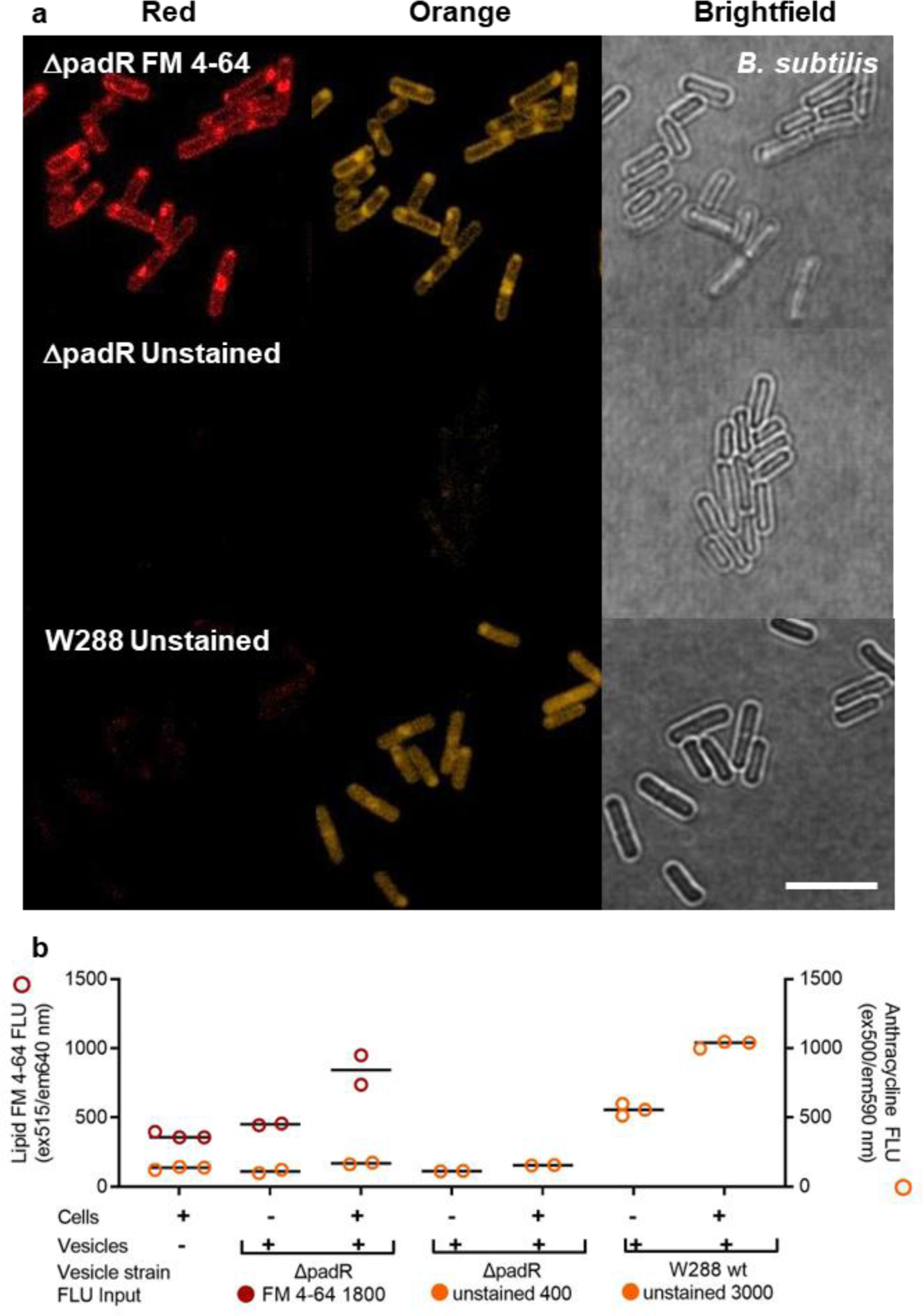
WAC00288 vesicles deliver fluorescent dyes and native cargo to membranes or cytoplasm of other microbial cells. *B. subtilis* incubated with vesicles from anthracycline knockout strain ΔpadR, equal amounts labeled with FM 4-64 (red circles) or unstained, or unstained (orange circles) vesicles from wildtype WAC00288 for 1 h. Cells were pelleted (5000xg, 5 min) and washed 100 fold with saline, then placed on an agarose pad and imaged by confocal microscopy for fluorescence (Red (ex515 nm, em650 – 760 nm), or Orange (ex488 nm, em570 – 650 nm)) and brightfield (T-PMT) (A). White bar, 5 µm. Conditions grouped in panels are from parallel treatments in one experiment, placed on one agarose pad, imaged and processed with the same settings. (B) Cells were also assayed for fluorescence with a spectrophotometer (open circles). Controls for expected background fluorescence were vesicles or cells alone (open circles). Fluorescence measured for FM 4-64 at ex 515, em 640 nm (red, left y-axis), and for anthracyclines at ex 500, em 590 nm (orange, right y-axis).

**Supplementary Figure 6.**
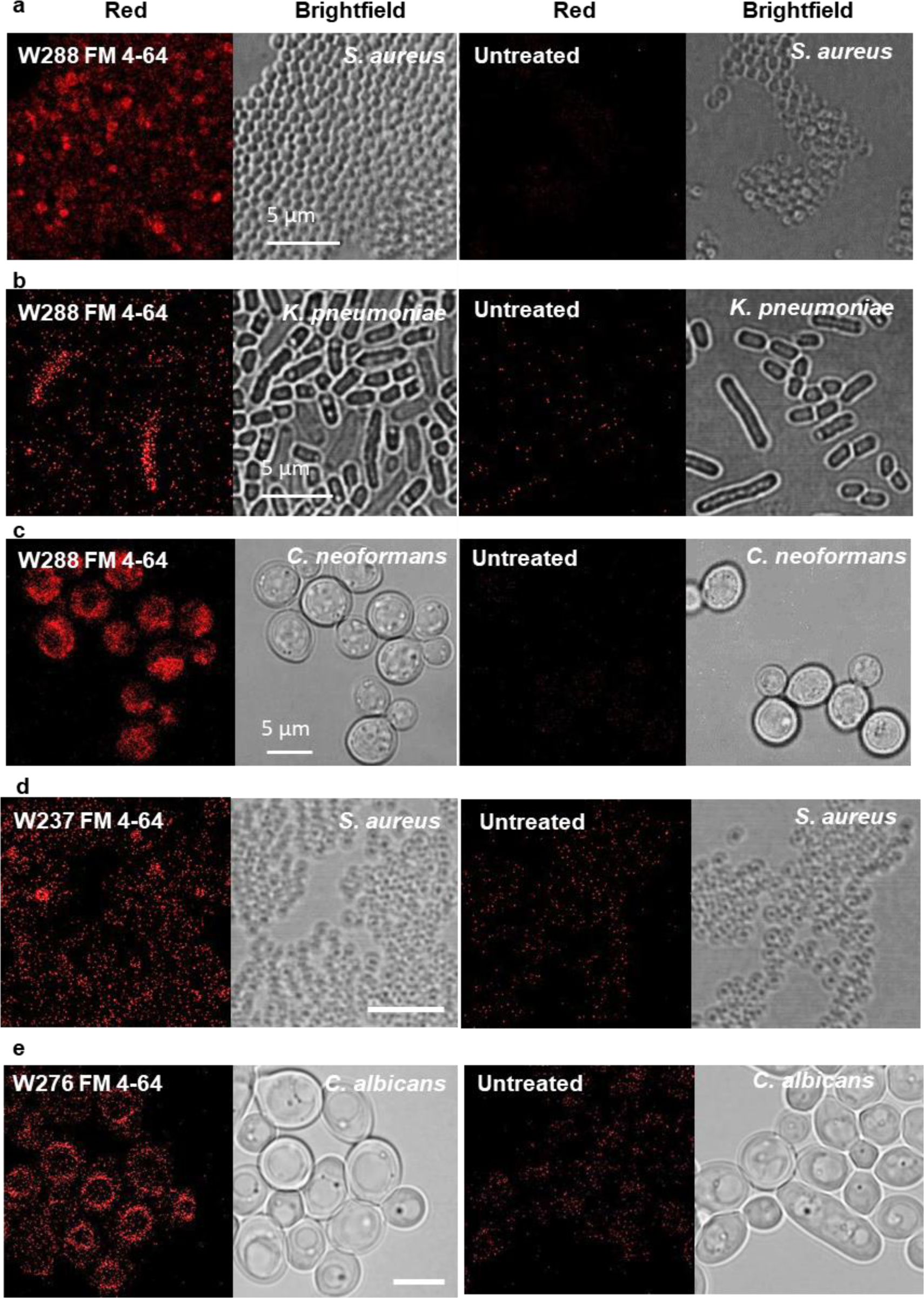
Transfer of fluorescence from Streptomyces vesicles to multiple microbial cell types. Cells were incubated with FM 4-64 labeled vesicles for 1 h, or left untreated, washed, and imaged in parallel under the same conditions for red fluorescence (ex488 nm, em650 – 760 nm). White bar 5 µm. (A-C) *S. aureus, K. pneumoniae*, or *C. neoformans* with FM 4-64 labeled WAC00288 vesicles. (D) S. aureus with FM 4-64 labeled WAC00237 vesicles. (E) C. albicans with FM 4-64 WAC00276 vesicles.

**Supplementary Table 1.** Antimicrobial activity and lipid fluorescence of high and low molecular weight fractions (HMW and LMW) from filamentous actinomycetes.

| Strain | Size fraction | Antimicrobial activity |  |  | Lipid signal |
| --- | --- | --- | --- | --- | --- |
|  | HMW >100 kDa<br>LMW <100 kDa | Gram-positive<br>(*S. aureus*) | Gram-negative<br>(*K. pneumoniae*) | Fungi<br>(*C. albicans*) | HMW – LMW<br>(FLU) |
| *S. aureofaciens* | HMW | - | - | - | 140 |
|  | LMW | + | + | - |  |
| *S. coelicolor* M145 | HMW | + | - | - | 1500 |
|  | LMW | + | - | - |  |
| *S. ghanaensis* | HMW | - | - | - | -390 |
|  | LMW | + | - | - |  |
| *S. griseus* | HMW | - | - | - | 4700 |
|  | LMW | ++ | + | - |  |
| *S. flavoperspicus* * | HMW | - | - | - | 2700 |
|  | LMW | + | + | - |  |
| *S. pristinaespiralis* | HMW | - | - | - | -140 |
|  | LMW | + | - | - |  |
| WAC00187 | HMW | + | - | - | 1700 |
|  | LMW | ++ | - | - |  |
| WAC00192 | HMW | - | - | - | 6700 |
|  | LMW | + | - | - |  |
| WAC00206 | HMW | - | - | - | 5200 |
|  | LMW | + | - | - |  |
| WAC00218 | HMW | + | - | - | 5500 |
|  | LMW | ++ | - | + |  |
| WAC00237 | HMW | + | - | - | 59000 |
|  | LMW | ++ | + | - |  |
| WAC00240 | HMW | + | - | - | 13000 |
|  | LMW | ++ | + | + |  |
| WAC00213 | HMW | - | - | - | 16000 |
|  | LMW | + | + | + |  |
| WAC00265 | HMW | - | - | ++ | 29000 |
|  | LMW | - | - | + |  |
| WAC00276 | HMW | - | - | ++ | 24000 |
|  | LMW | - | - | + |  |
| WAC00288 | HMW | ++ | - | - | 16000 |
|  | LMW | + | - | - |  |
| WAC00303 | HMW | + | - | + | 16000 |
|  | LMW | - | - | - |  |

\*genetically modified strain to express cryptic gene clusters<sup>54</sup>

